# The role of Gdf5 in the development of the zebrafish fin endoskeleton

**DOI:** 10.1101/2021.01.31.428959

**Authors:** Laura Waldmann, Jake Leyhr, Hanqing Zhang, Amin Allalou, Caroline Öhman-Mägi, Tatjana Haitina

## Abstract

**Background:** The development of the vertebrate limb skeleton requires a complex interaction of multiple factors to facilitate correct shaping and positioning of bones and joints. Growth and differentiation factor 5 (Gdf5), a member of the transforming growth factor-beta family (TGF-β) is involved in patterning appendicular skeletal elements including joints. Expression of *gdf5* in zebrafish has been detected within the first pharyngeal arch jaw joint, fin mesenchyme condensations and segmentation zones in median fins, however little is known about the functional role of Gdf5 outside of Amniota.

**Results:** We generated CRISPR/Cas9 knockout of *gdf5* in zebrafish and analysed the resulting phenotype at different developmental stages. Homozygous *gdf5* mutant zebrafish displayed changes in segmentation of the endoskeletal disc and, in consequence, loss of posterior radials in the pectoral fins. Mutant fish also displayed affected organisation and length of skeletal elements in the median fins, however joint formation and mineralisation process seemed unaffected.

**Conclusions:** Our study demonstrates the importance of Gdf5 for the paired and median fin endoskeleton development in zebrafish and reveals that the severity of the effect increases from anterior to posterior side of the elements. Our findings are consistent with phenotypes observed in human and mouse appendicular skeleton in response to *Gdf5* knockout, suggesting a broadly conserved role for Gdf5 in Osteichthyes.

## 1 INTRODUCTION

Skeletal development requires a complex interaction of multiple factors to ensure correct positioning and shaping of cartilage elements and bones including proper formation of joints articulating the skeleton. Skeletogenesis via endochondral ossification involves three major processes starting with mesenchyme condensation followed by chondrogenesis and finally the invasion of blood vessels and chondrocyte replacement by and transformation into osteoblasts (Aghajanian and Mohan, 2018; Mackie et al., 2008). Members of the Transforming Growth Factor beta (TGF-β) ligand superfamily including Bone Morphogenic Proteins (BMPs) and Growth Differentiation Factors (GDFs) play a major role during all stages of skeletogenesis (Salazar et al., 2016). GDF5, also called BMP14, CDMP1, and Contact, has been shown to be involved in the development of the appendicular skeleton (Storm et al., 1994; Storm and Kingsley, 1999).

Gdf5, like other BMPs, is initially produced as a large precursor protein containing an N-terminal signal peptide, a prodomain, and a C-terminal bioactive mature domain. A small cleavage site connects the prodomain and mature domain, and the mature domain is cleaved away from the rest of the precursor protein to form the active ligand (Katagiri and Watabe, 2016). The Gdf5 ligand binds to type I and type II serine-threonine kinase receptors such as BMPR-IB (Baur et al., 2000; Klammert et al., 2015; Nishitoh et al., 1996) and triggers intracellular signaling pathways that regulate expression of target genes.

Expression of *Gdf5* in amniotes is largely restricted to limbs, whereas amphibians and ray-finned fishes display expression in the head in addition to the appendicular skeleton (Bruneau et al., 1997; Miller et al., 2003; Schwend and Ahlgren, 2009; Square et al., 2015). The expression pattern of *Gdf5* in amniotes is known from studies in chicken and mouse models. *Gdf5* expression is first detected during early stages of limb skeletogenesis in the knee and elbow joints (Shwartz et al., 2016), and in the mesenchyme condensations of the digital rays in the autopod at the positions where the articulations between metacarpals and proximal phalanges will develop (Merino et al., 1999; Storm and Kingsley, 1996). At the onset of chondrogenesis, *Gdf5* expression in the metatarsophalangeal, metacarpophalangeal, and interphalangeal joints is upregulated. As joint maturation continues and cavitation takes place, *Gdf5* is downregulated and primarily restricted to lateral joint edges (Merino et al., 1999). Similarly, in the appendicular skeleton of *Xenopus, gdf5* is first expressed in the limb bud and then restricted to joint forming regions during later stages in the autopod (Satoh et al., 2005). Craniofacial *gdf5* expression in *Xenopus* is found in the developing intramandibular joints before skeletal differentiation (Square et al., 2015).

In the teleost model zebrafish, *gdf5* is expressed in the pharyngeal arch primary jaw joint and at the midline of the future basihyal cartilage element (Miller et al., 2003; Schwend and Ahlgren, 2009). In the pectoral fins, *gdf5* is first expressed in the fin bud mesenchyme in the antero-proximal region and as development continues into the pec-fin stage, expression expands posteriorly along the proximal margin of the fin bud (Bruneau et al., 1997). The expression pattern of *gdf5* in zebrafish pectoral fins beyond 90 hpf has not been reported, consequently little is known about the role of *gdf5* in the later stages of pectoral fin development, which includes the segmentation of the endoskeletal disc into the four proximal radials, and the condensation of the distal radials.

In the dorsal and anal fin of zebrafish, *gdf5* is first expressed in the mesenchyme between the prefigured radials and at later stages within the segmentation zone when the distal radials segment away from the proximal radials (Crotwell et al., 2001). *gdf5* expression in the caudal fin begins in the mesenchyme anterior to the parhypural and hypural 1, and as hypurals 2-5 start to form, expression can be detected between those condensations. A small population of gdf5-positive cells is furthermore present posterior to the epurals (Crotwell et al., 2001).

As with other members of the BMP family, Gdf5 has been shown to regulate chondrogenesis in a temporo-spatial manner (Storm and Kingsley, 1999; Tsumaki et al., 1999).

Overexpression of Gdf5 at the mesenchyme condensation stage in chick and mouse autopods results in enhanced recruitment of mesenchymal cells (Tsumaki et al., 1999;

Buxton et al., 2001). It has furthermore been shown that the supply of exogenous GDF5 protein leads to an increased commitment of chondrocytes to undergo hypertrophy in both chick and mouse limbs (Buxton et al., 2001; Coleman and Tuan, 2003; Storm and Kingsley, 1999). As a consequence of enhanced differentiation and proliferation of chondroprogenitor cells and chondrocytes, the skeletal elements of the autopods increase in size. During joint development in mice and chicken autopods, *Gdf5* is involved in restricting joint markers to the joint formation site (Storm and Kingsley, 1999). Beyond early joint-forming processes, a continuous influx of Gdf5-positive cells is involved in the differentiation of various synovial joint tissues including the articular cartilage and associated ligaments (Shwartz et al., 2016).

Several disease and mutant phenotypes are associated with *GDF5* mutations. The intensively-studied mouse *brachypodism* phenotype is caused by a frameshift mutation within *Gdf5* resulting in a premature stop codon (Storm et al., 1994). The murine *brachypodism* phenotype is characterised by reduced length of limb long bones and deformations in the digits including loss of joints, size reduced phalanges and loss of skeletal elements (Gruneberg and Lee, 1973; Storm and Kingsley, 1996). Similar phenotypic effects can be seen in human autosomal recessive syndromes Grebe type and Hunter-Thompson, also linked to mutations in *GDF5* (Thomas et al., 1996; Martinez-Garcia et al., 2016).

In contrast, the effects of *GDF5* mutation or loss in non-mammalian species have not been studied until now. In this study we generated *gdf5* knockout zebrafish line with CRISPR/Cas9 genome editing and analysed the resulting skeletal defects in larval and adult fish. We demonstrate that homozygous *gdf5* mutant zebrafish display truncated median fin endoskeletal elements and loss of posterior radials in the pectoral fins.

## 2 RESULTS

### 2.1 Generation of *gdf5*^uu3703/uu3703^ knockout line

By using CRISPR-Cas9 genome editing approach in zebrafish we introduced a 5bp deletion in exon 2 of the *gdf5* gene. This deletion induces a frameshift that results in a premature stop codon, truncating the produced protein from 474 amino acids (aa) to 230 aa (Figure 1A, B). This truncated protein is missing a large portion of the prodomain and the entirety of the cleavage site and mature domain. Heterozygous *gdf5*^+/uu3703^ zebrafish were incrossed to generate homozygous mutants. Both *gdf5*^+/uu3703^ and *gdf5*^uu3703/uu3703^ fish displayed no outward morphological alterations relative to wild-type siblings (Figure 1C).

**FIGURE 1.**
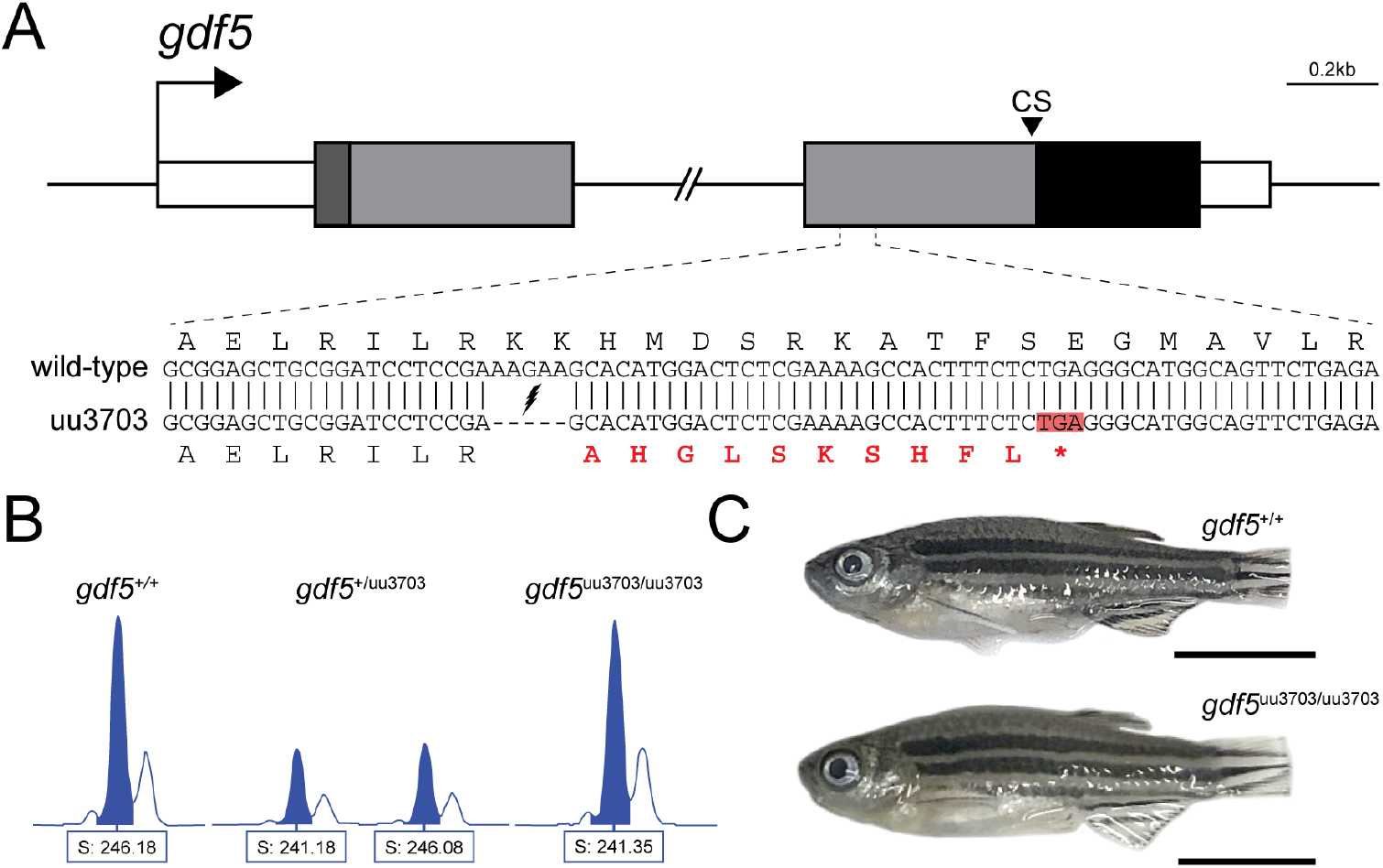
Zebrafish *gdf5* knockout generated with CRISPR/Cas9. **(A)** To-scale schematic of the two-exon *gdf5* gene locus on chromosome 6, with the 4,267bp intron truncated. White boxes represent the 5’ and 3’ UTRs, the dark grey box represents the sequence coding for the 22 aa signal peptide, the light grey boxes represent the sequences coding for the rest of the prodomain (334 aa), and the black box represents the sequence coding for the 118 aa mature domain. “CS” indicates the position of the 5 aa cleavage site. The sequence of the zoomed in section of exon 2 shows the alignment between the wild-type and mutant (uu3703) alleles. The mutant allele has a 5-base deletion that results in a frameshift and a premature stop codon highlighted in red. **(B)** Wild-type, heterozygous *gdf5*^+/uu3703^, and homozygous *gdf5*^uu3703/uu3703^ fish were identified by fragment length analysis. The wild-type displayed one peak (246.18), *gdf5*^+/uu3703^ displayed one wild-type peak (246.08) and one mutant peak (241.18), and *gdf5*^uu3703/uu3703^ displayed one mutant peak (241.35). **(C)** At 60 dpf there is no outward phenotypic difference between *gdf5*^uu3703/uu3703^ and wild-type (caudal fins were clipped for genotyping). Scale bars: 5 mm.

### 2.2 The development of the pectoral fin skeleton is affected in *gdf5*^uu3703/uu3703^ fish

We examined a range of developmental stages between larval and adult zebrafish using bone and cartilage staining and μCT to assess the effects of *gdf5* knockout on skeletal structures. Comparison to wild-type zebrafish revealed that *gdf5*^uu3703/uu3703^ fish displayed the typical stages of pectoral fin development up to and including the establishment of the endoskeletal disc cartilage. However, *gdf5*^uu3703/uu3703^ diverged from *gdf5^+/+^* after the formation of the first cartilage subdivision zone (CSZ1) at 30 dpf (7.0mm standard length (SL)). CSZ1 appears significantly larger in *gdf5* mutants (Figure 2A-B). At 7.2mm SL wild-type fish established CSZs 2 and 3, subdividing two cartilage condensations into four, corresponding to proximal radials 1-4 (Figure 2C). At the same SL, *gdf5*^uu3703/uu3703^ fish appeared to either expand CSZ1, or merge CSZs 1-3 into a single large central subdivision zone, leaving two elements similar in position and shape to the cartilaginous anlage of wild-type proximal radials 1 and 4, while radials 2 and 3 were absent (Figure 2D).

**FIGURE 2.**
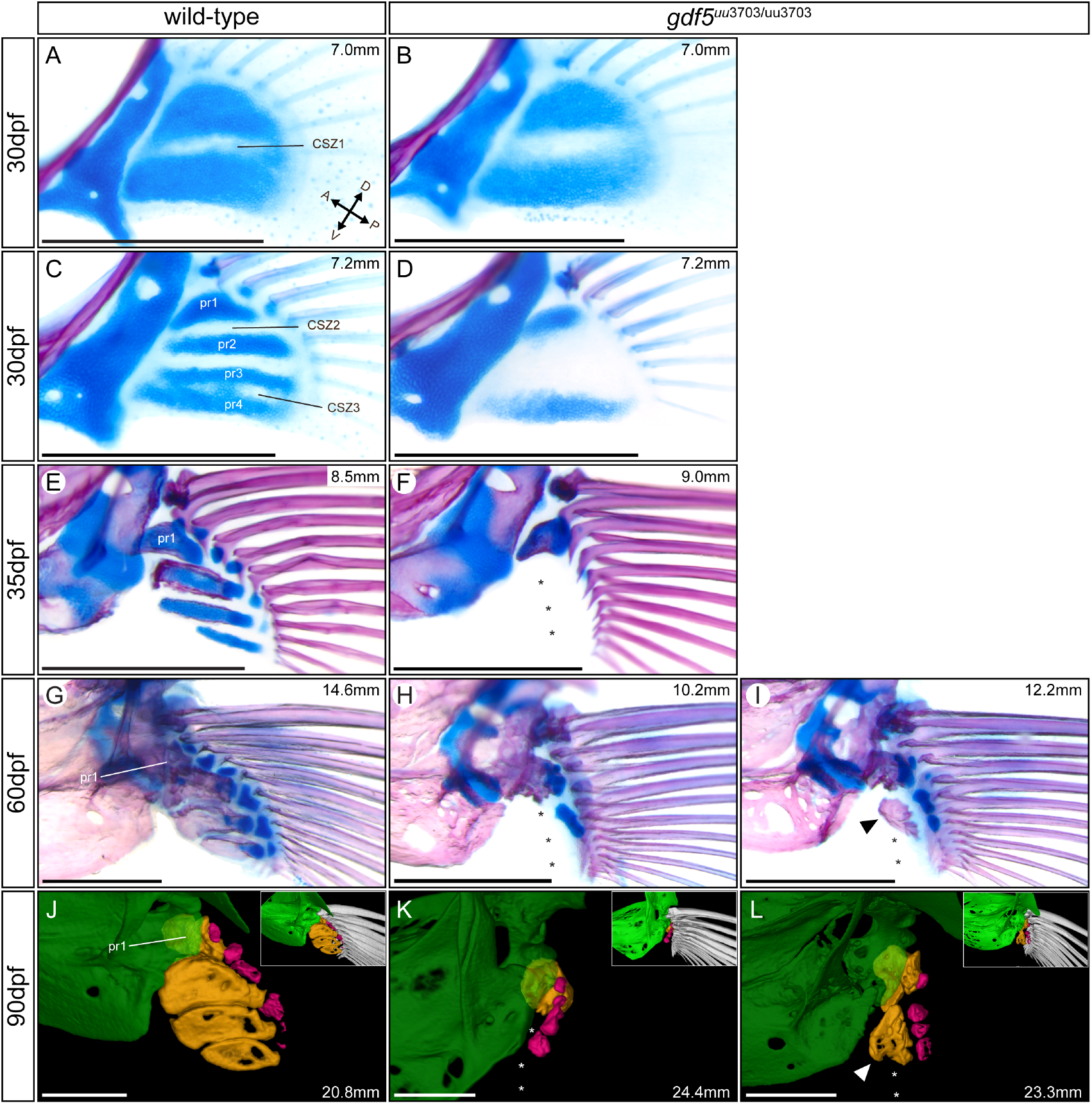
*gdf5*^uu3703/uu3703^ display loss of pectoral fin radials. All images depict pectoral fins with postcleithra removed in a dorsomedial view. Bone- and cartilage-stained pectoral fins of wild-type and *gdf5*^uu3703/uu3703^ zebrafish at 30 dpf **(A, B, C, D**) and 35 dpf **(E, F)**, respectively. **(G)** pectoral fin of 60dpf wild-type zebrafish. **(H, I)** pectoral fins of two *gdf5*^uu3703/uu3703^ zebrafish siblings displaying variable presence of bone at the position of proximal radial 2. **(J)** pectoral fin of 90 dpf wild-type zebrafish following μCT scanning and 3D segmentation. **(K, L)** pectoral fin of 90 dpf *gdf5*^uu3703/uu3703^ zebrafish following μCT scanning and 3D segmentation. The measurements in mm refer to standard length (SL). Asterisks highlight the absence of proximal radials 2-4. Arrowheads indicate the variably present bone in place of proximal radial 2 in *gdf5*^uu3703/uu3703^ fish. In μCT images, the pectoral girdle (green) is rendered partially transparent to provide unobstructed views of the proximal radials (orange). Fin rays (white) were removed for clarity and shown instead in the insets. Distal radials are coloured in pink. pr - proximal radial; CSZ - cartilage segmentation zone. Scale bars: 500 μm.

At 30 dpf chondrocytes in the proximal radial condensations of both wild-type and *gdf5* mutant fish appeared to undergo mitotic divisions apparent by thinner lines dividing the cells (Figure 2A-B). However, the total number of chondrocytes in forming radials looked significantly reduced in *gdf5* mutant fish at 7.2mm SL. By 35 dpf, wild-type pectoral fins displayed four proximal radials beginning the process of endochondral ossification, while all *gdf5* mutant pectoral fins possessed just proximal radial 1 (reduced in size), and were entirely missing proximal radials 2-4 (Figure 2E-F).

In approximately half of all examined *gdf5*^uu3703/uu3703^ pectoral fins, this phenotype persisted until adulthood (60-90 dpf), while in the other half a second bony element became evident by 60dpf in the position normally occupied by proximal radial 2, although it is smaller and misshapen compared to this element in wild-type zebrafish (Figure 2G-I). We were unable to observe the origin of this bone but based on the fact that we never found it in association with any cartilage, and no potential cartilaginous precursor elements were observed in earlier developmental stages, we infer that it may not arise via endochondral ossification, and as such is developmentally distinct from wild-type proximal radial 2. We observed several adult zebrafish that possessed this bony element in only one of paired pectoral fins, suggesting it may be a transient ectopic element dependent on plastic local signaling.

A similar phenotype was observed in the distal radials that articulate the proximal radials with the lepidotrichia. The posterior distal radials, those that would be associated with proximal radials 3 and 4, were absent while the more anterior distal radials were present, albeit reduced at 30-60 dpf relative to wild-type (Figure 2F, H, I). By 90dpf, these anterior distal radials appeared to have ossified to a similar extent as wild-type distal radials (Figure 2J-L). The shoulder girdle and pectoral lepidotrichia appeared to develop normally in all examined *gdf5*^uu3703/uu3703^ fish.

### 2.3 The organisation and length of skeletal elements in median fins is affected in *gdf5*^uu3703/uu3703^ fish

To investigate the skeletal phenotypes of *gdf5*^uu3703/uu3703^ fish we examined median fins in larvae, juveniles and adults by skeletal staining and μCT. At 14 dpf, when cartilage condensations of the caudal fin begin to form, a prominent phenotype could be detected in *gdf5*^uu3703/uu3703^ fish. At this stage, the parhypural and hypural 1 displayed a separation within their respective developing cartilage condensations in *gdf5*^uu3703/uu3703^ fish (Figure 3A, A’, B, B’). The other hypurals however were not drastically affected at this timepoint (Figure 3A, A’, B, B’). All the skeletal elements of the caudal fin were present in wild-type fish by 30 dpf and underwent partial endochondral ossification (Figure 3C, C’’). At this time point, hypurals 3-5 of *gdf5*^uu3703/uu3703^ fish were severely truncated while the more ventrally located elements (hemal spines, parhypural, hypurals 1-2 were only slightly deformed (Figure 3D, D’’). μCT analysis of mutant fish at 90 dpf showed the same trend whereby all the hypurals and the parhypural were truncated, but hypurals 3-5 were the most severely deformed (Figure 4C, F). Ossification of the hypurals appeared to proceed in *gdf5*^uu3703/uu3703^ fish as in wild-type (Figure 3C-D; 4C, F).

**FIGURE 3.**
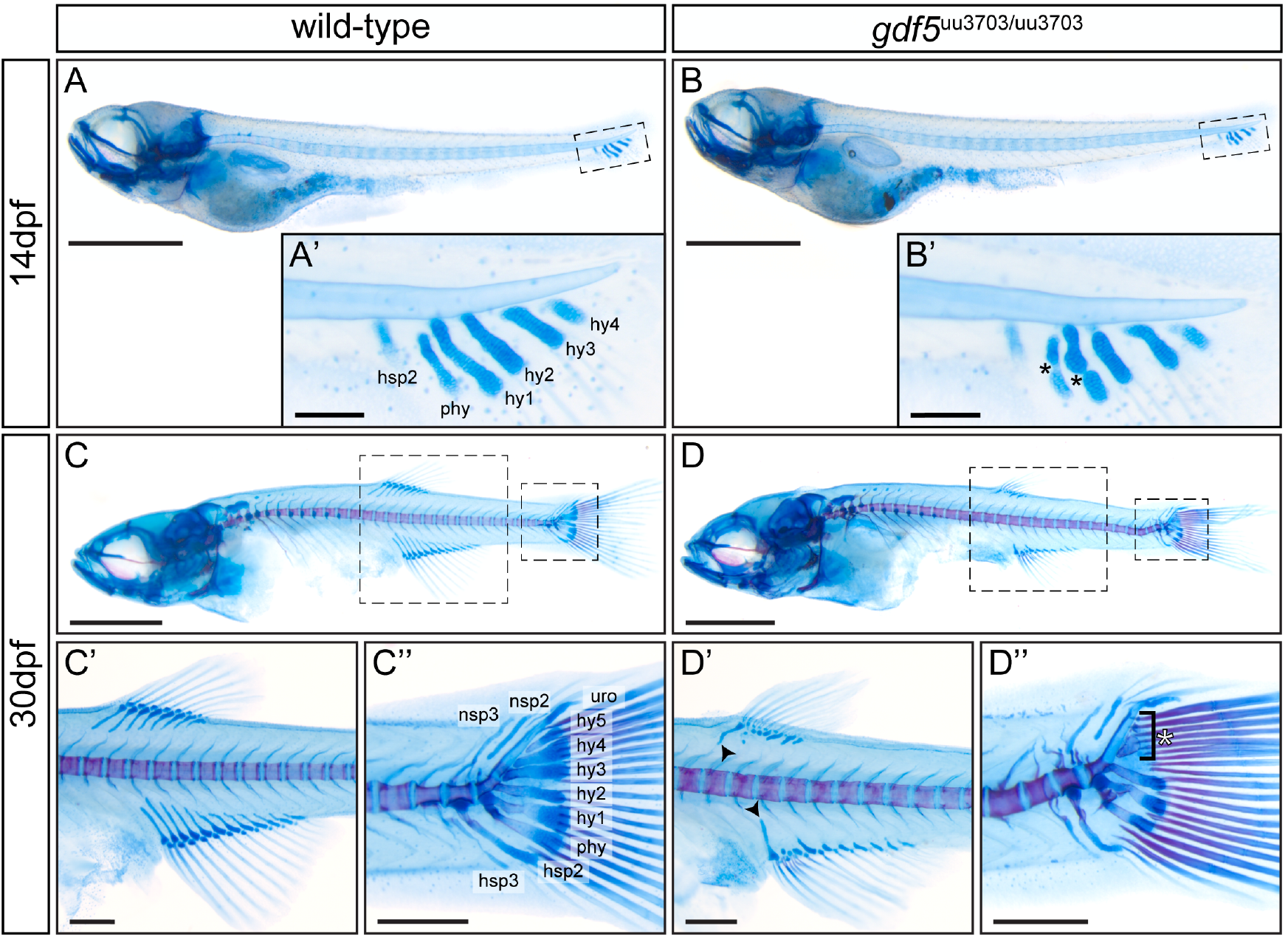
Skeletal staining reveals abnormalities in median fin skeletal organization in *gdf5*^uu3703/uu3703^ zebrafish. Lateral views of cartilage- and bone-stained wild-type fish at 14 dpf **(A, A’)** and 30 dpf **(C, C’, C’’)**; and *gdf5*^uu3703/uu3703^ fish at 14 dpf **(B, B’)** and 30 dpf **(D, D’, D’’)**. Dashed boxes **(A-D)** mark magnified regions **(A’-D’’). (B’)** *gdf5*^uu3703/uu3703^ larvae display separation within the parhypural and hypural 1 cartilage condensations (black asterisks) **(D’)** Dorsal and anal fin radials are truncated in *gdf5*^uu3703/uu3703^ fish. The most anterior proximal radials are less affected in both dorsal and anal fin (black arrowhead). **(D’’)** *gdf5*^uu3703/uu3703^ fish display truncated and slightly deformed and truncated hemal spines, parhypural and hypural 1. Hypural 3-5 are severely shortened in size (white asterisk). hsp - hemal spine, phy - parhypural, hy - hypural, uro - uroneural, ep - epural, ns - neural spines. Scale bars: 1 cm **(A-D)**, 100 μm **(A’ and B’)** and 250 μm **(C’-D’’)**.

**FIGURE 4.**
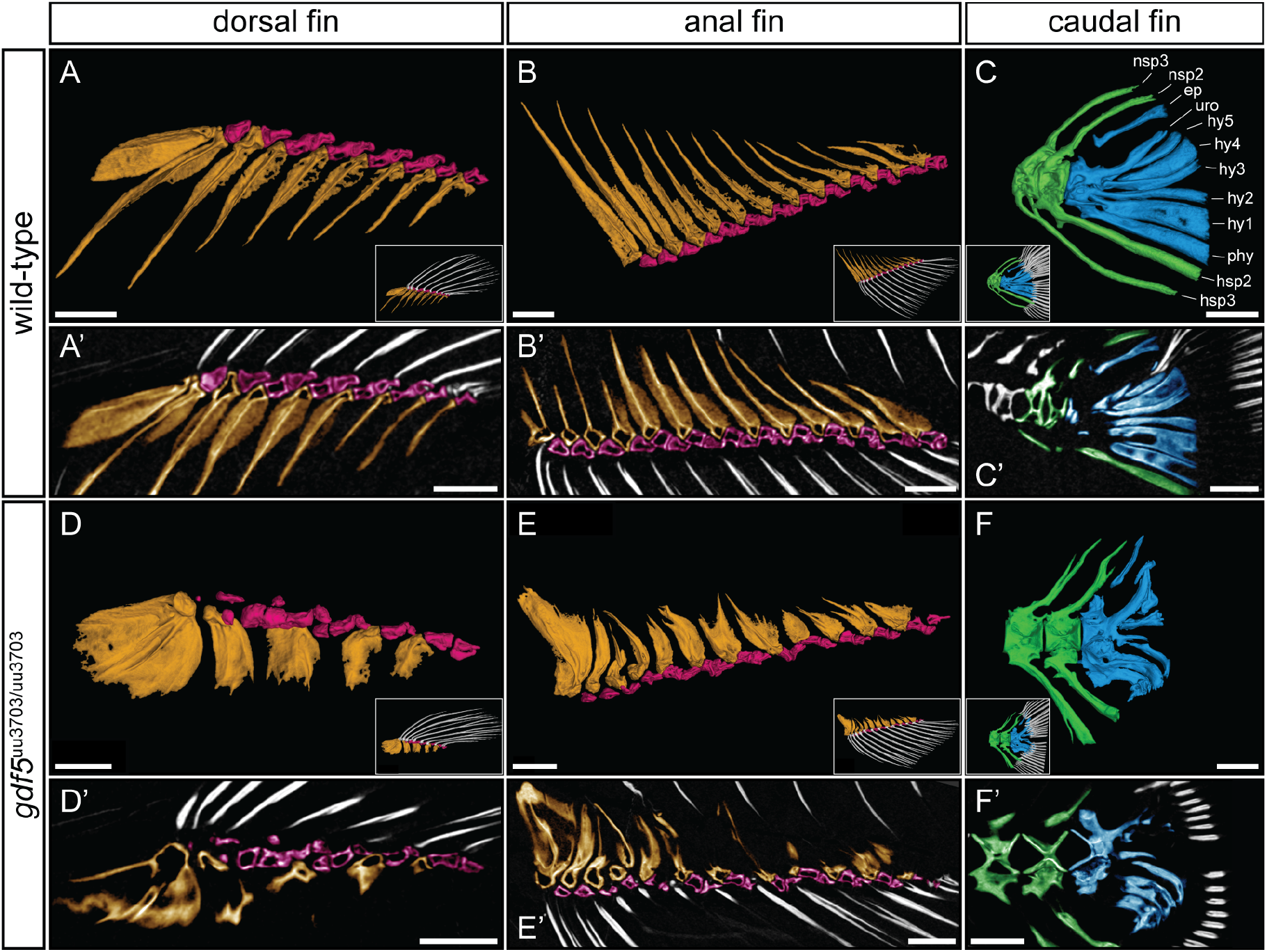
*gdf5*^uu3703/uu3703^ display defects in median fin skeletal elements. At 90 dpf wild-type **(A-C)**, and *gdf5*^uu3703/uu3703^ **(D-F)** lateral views of the dorsal, anal, and caudal fin skeleton following μCT scanning and 3D segmentation (anterior to left). Fin rays (white) were removed for clarity and shown instead in the insets. **(A’-F’)** False-coloured virtual thin sections of median fin skeletal elements. **(A’, B’, D’, E’)** 20 μm sections, **(C’, F’)** 30 μm sections. Proximal radials are coloured in orange, distal radials in pink, epural, hypurals, and parhypural in blue, and preurals in green. ep - epural; hsp - hemal spine; hy - hypural; nsp neural spine; phy - parhypural; uro - urostyle. Scale bars: 500 μm.

At 30 dpf, when both the dorsal and anal fin are well-developed, *gdf5*^uu3703/uu3703^ fish displayed a significant disorganization of the proximal and distal radials compared to wild-type fish (Figure 3C’, D’). The anterior-most proximal radial in both the dorsal and anal fin was typically less affected in *gdf5*^uu3703/uu3703^ fish and often projected towards the vertebral column as in wild-type (Figure 3D’). In contrast, the other proximal radials were variably truncated and often barely formed any cartilage outgrowth towards the vertebrae. Distal radials in mutant fish were present in both dorsal and anal fin but appeared less developed at this stage (Figure 3C’, D’). μCT analysis of dorsal and anal fin in 90 dpf *gdf5*^uu3703/uu3703^ fish was consistent with observations of skeletal-stained 30 dpf fish (Figure 4A, B, D, E). Proximal radials were extended in width and did not show any proximal cartilage outgrowth, however appeared to ossify normally (Figure 4D, D’, E, E’). Lepidotrichia in all median fins were present and not affected in either growth or ossification in *gdf5*^uu3703/uu3703^ fish.

### 2.4 *gdf5*^uu3703/uu3703^ fish display unaffected joint formation and cartilage maturation in median fin skeletal elements

*gdf5* is expressed during early radial cartilage condensation and within the segmentation region when the distal tip of each condensation segments away to form the distal radial in both the dorsal and anal fin (Crotwell et al., 2001). To investigate whether Gdf5 deficiency had an effect on the joint, which articulates the proximal and distal radials we performed histological staining of adult wild-type and mutant thin sections. Histological sections were furthermore used to analyse whether Gdf5 deficiency affected cartilage maturation in median fin skeletal elements.

We analysed 60 dpf wild-type and *gdf5*^uu3703/uu3703^ HE stained thin sections of anal, dorsal, and caudal fins (Figure 5). Dorsal and anal fin articulations between the proximal and distal radials displayed a layer of joint-characteristic cells at the articulating surfaces of the elements (Figure 5A’-D’). In contrast to hypertrophic chondrocytes present within the radials in both wild-type and *gdf5*^uu3703/uu3703^ fish, these cells appeared similar to perichondrium-forming cells (Figure 5A’-D’). We could not detect any proximal-distal joint defects in *gdf5*^uu3703/uu3703^ fish (Figure 5B’, D’). The joints displayed a cavity, which seemed to be enclosed by a capsule and appeared unaffected in *gdf5* mutant fish when compared to wild-type (Figure 5A’-D’). No apparent delay in chondrocyte maturation could be detected in *gdf5*^uu3703/uu3703^ fish median fin skeletal elements when compared to wild-type (Figure 5A’-F’).

**FIGURE 5.**
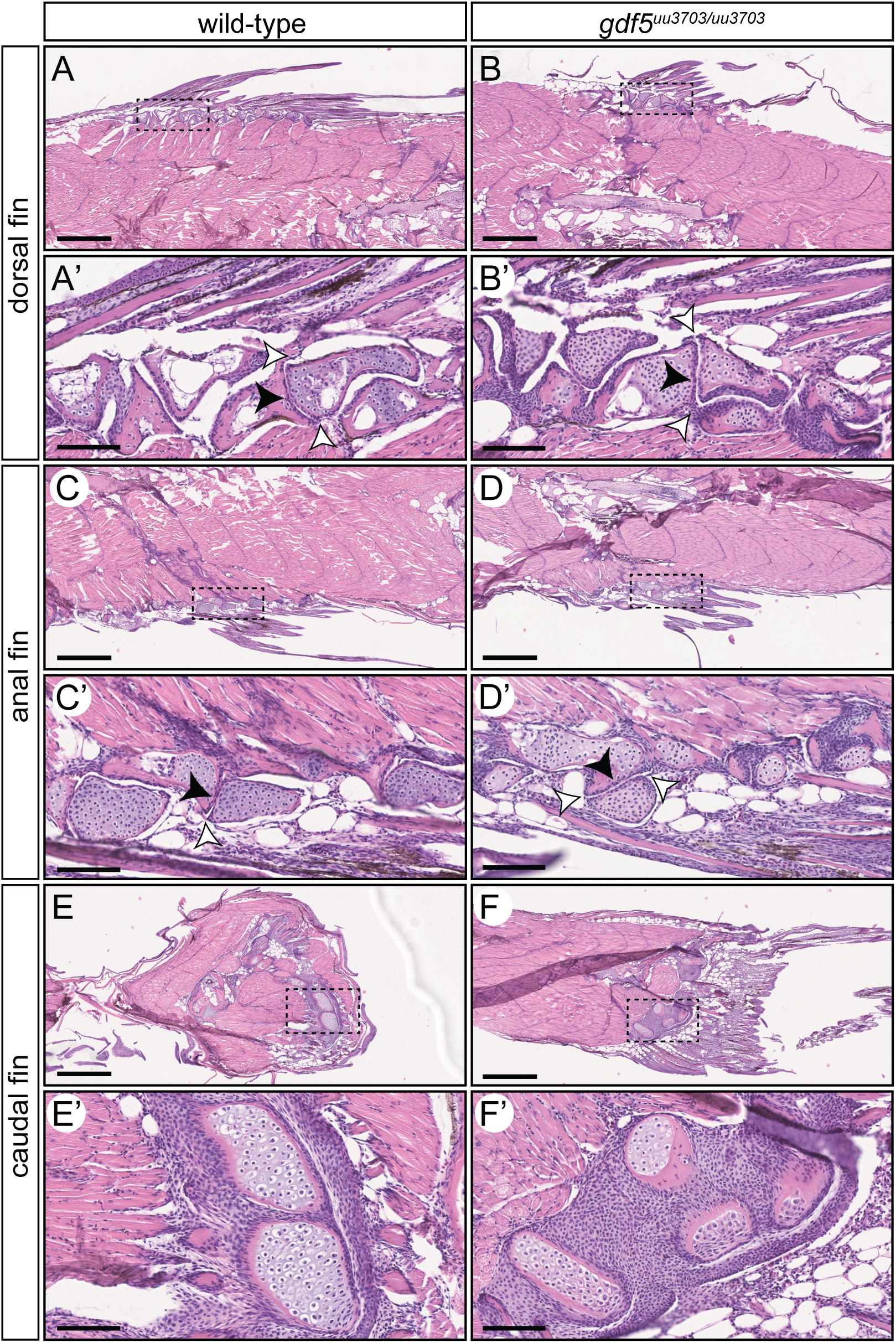
Median fin histology of adult wild-type and *gdf5*^uu3703/uu3703^ zebrafish reveals intact joints between distal and proximal radials and normal maturation of the skeletal elements. Sagittal sections from 60dpf of wild-type **(A, A’, C, C’, E, E’)** and *gdf5* mutant **(B, B’, D, D’, F, F’)** zebrafish median fins stained with Hematoxylin and Eosin (HE). Anterior to the left, dorsal to the top. Dashed boxes in **A-F** mark magnified region in **A’-F’**. **(A, A’, B, B’)** Dorsal fin; **(C, C’, D, D’)** anal fin; **(E, E’, F, F’)** caudal fin. (B’, D’) Black arrowheads indicate joint capsule, white arrowheads indicate articular cartilage. All joints are intact and do not display any differences when compared to wild-type (A’, C’). (F, F’) Caudal fin skeletal elements display no apparent delays in chondrocyte maturation when compared to wild-type (E, E’). Scale bar: 500 μm **(A-F),** 100 μm **(A’-F’)**.

### 2.5 *gdf5*^uu3703/uu3703^ fish display unaffected head skeleton

OPT analysis of cartilage- and bone-stained 9 dpf zebrafish larvae revealed no skeletal abnormalities in the skull in *gdf5* mutants compared to wild-type (Figure 6). The cartilage of Meckel’s cartilage and the palatoquadrate appeared to be unaffected, including the shape of the jaw joint (Figure 6D, H, Supplemental Video 1). The only observable difference was reduced ossification of the anterior notochord in the *gdf5*^uu3703/uu3703^ mutants relative to wild-types (Figure 6E, F). No differences in the ossification of the notochord were observed at older ages and there was a slight (~4%) reduction in head length in the 9 dpf mutants relative to wild-types, suggesting that this difference in anterior notochord ossification might be explained by a general growth phenotype rather than a notochord-specific phenotype. To further investigate whether *gdf5*^uu3703/uu3703^ displayed reduced body size compared to siblings the standard length (SL) of a large sample of mutants and wild-type zebrafish between 28 and 34 dpf were measured and compared (Supplemental Figure 1). No significant difference in SL was found, and the SLs of both groups were consistent with normal growth curves (Parichy et al., 2009), suggesting that by this juvenile stage of development, any potential growth phenotype was too insignificant to be detected, or that there was in fact no growth phenotype at all, and the slight difference in head size at 9 dpf was due to random chance.

**FIGURE 6.**
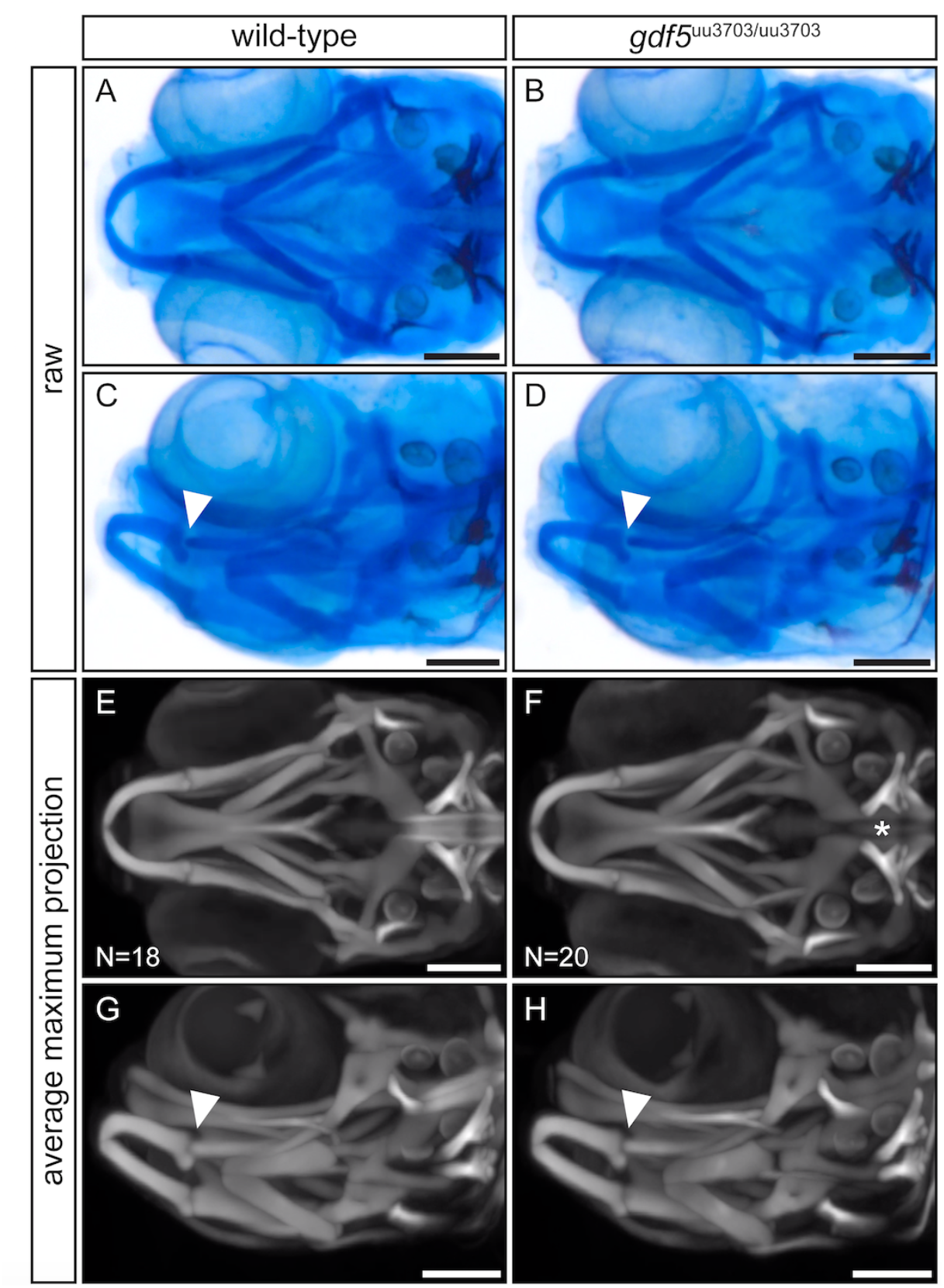
Optical Projection Tomography reveals no craniofacial skeletal defects in *gdf5*^uu3703/uu3703^ zebrafish at 9 dpf. **(A-D)** Examples of raw images of single 9 dpf zebrafish heads captured during OPT. **(E-H)** Maximum projection views of averaged 3D models generated by OPT analysis of 18 wild-type and 20 *gdf5* mutant zebrafish at 9 dpf. Arrowheads indicate the jaw joint between Meckel’s cartilage and the palatoquadrate. The asterisk highlights the reduced anterior notochord ossification in *gdf5*^uu3703/uu3703^ fish relative to wild-types. Scale bars: 150μm.

## 3 DISCUSSION

The role of Gdf5 in skeletal development has been extensively studied in amniote model species while little is known about its function in other vertebrate groups. Here we present the first phenotypic analysis of homozygous *gdf5* mutant zebrafish *(Danio rerio)* at multiple developmental stages. Our results show skeletogenic defects in pectoral and median fin development demonstrating a largely conserved role for Gdf5 in skeletal developmental while also highlighting divergent functions that may be specific to actinopterygians.

The pectoral fin buds of *gdf5*^uu3703/uu3703^ zebrafish appear morphologically normal until segmentation of the radials takes place, suggesting that the early A-P patterning is not affected by the Gdf5 deficiency. On the other hand, given that the anterior proximal radial is largely unaffected in *gdf5*^uu3703/uu3703^ fish and the posterior distal radials, which condense independently of the posterior proximal radials, are lost, there is clearly a posterior-specific effect on the pectoral endoskeleton. This could suggest that there are early effects on A-P patterning that manifest much later in development, or that Gdf5 action is essential specifically during the period of development around segmentation and in posterior pectoral tissues. Little is currently known about the genetic control of endoskeletal disc segmentation, so further research will be required to fully elucidate the role of Gdf5 in this process.

The pectoral fin of zebrafish is homologous to the mammalian forelimb, sharing skeletal elements that ossify endochondrally from cartilaginous anlage (Gehrke et al., 2015; Grandel and Schulte-Merker, 1998). However, the initial establishment of cartilaginous elements in the pectoral fin of actinopterygian fish is achieved quite differently compared to tetrapod limbs. In tetrapods, the endoskeleton is patterned prior to the condensation of mesenchyme into chondrogenic tissue, such that the condensations form in the pattern of skeletal elements directly (Shimizu et al., 2007). In contrast, teleosts initially form the endoskeletal disc before matrix decomposition and chondrocyte dedifferentiation subdivides the disc into a number of proximal radial elements (Dewit et al., 2011). This mechanism for the formation of proximal radials is broadly consistent in actinopterygians as a group, although the most basal extant species display a mosaic of developmental mechanisms in other cartilaginous elements of the pectoral fin (Davis et al., 2004; Grandel and Schulte-Merker, 1998). These differences make it difficult to assign homologies between tetrapod and teleost pectoral skeletal elements and to draw connections between mutant phenotypes.

*Gdf5* inactivation has been intensively studied in *brachypodism* mice and is responsible for human syndromes Hunter-Thompson- and Grebe-type chondrodysplasia (Martinez-Garcia et al., 2016; Storm and Kingsley, 1996; Thomas et al., 1996). The phenotypic effects caused by Gdf5 deficiency are restricted to the appendicular skeleton and lead to shortening of the skeletal elements. Here we show that *gdf5* inactivating mutation in zebrafish affects endochondrally ossifying skeletal elements of the median fins which are truncated and shortened in size, similar to what is seen in *Gdf5-*deficient mice and human. All radials in the dorsal and anal fin and the parhypural and hypurals in the caudal fin are affected in homozygous *gdf5* mutant zebrafish. Interestingly, more anteriorly located radials in the anal and dorsal fin and the more ventrally located parhypural and hypurals of the caudal fin are generally less affected by Gdf5 loss than the more posterior and dorsal elements in these structures, respectively. Ventrally and dorsally located elements of the caudal fin are anterior and posterior, respectively, in relation to the zebrafish body axis. This is also in agreement with anterior weak, posterior strong effect we observed in pectoral fin radials in *gdf5* mutant zebrafish (Figure 2).

Gdf5 has been shown to affect skeletogenesis during early stages by recruiting cells to the mesenchyme condensation and later stages within the joint interzone signaling into the epiphysis of the articulating elements. Overexpression studies led to increased skeletal element size in response to increased cell number of the prechondrogenic condensation (Francis-West et al., 1999a), and mesenchyme condensations in *brachypodism* mice were reduced in size due to a decreased cell number (Gruneberg and Lee, 1973). We have not analysed mesenchyme condensations in *gdf5* deficient zebrafish, however during the endoskeletal disc segmentation the total number of chondrocytes forming the radials appeared dramatically reduced (Figure 2D). Since in wild-type zebrafish *gdf5* expression is present in all affected skeletal elements and *gdf5* mutant fish displayed similar phenotypic characteristics as in *brachypodism* mice, we suggest that similar mechanism leads to the size reduction of the skeletal elements in both species.

Previous studies reported that Gdf5 promotes differentiation of mesenchymal cells into chondrocytes and chondrocyte maturation (Bai et al., 2004; Buxton et al., 2001; Coleman and Tuan, 2003; P. H. Francis-West et al., 1999a; Merino et al., 1999; Storm and Kingsley, 1999). This role of Gdf5 as a promoter of chondrocyte maturation is opposite to the function of NK 3 homeobox 2 (Nkx3.2) transcription factor, which is a well-known chondrocyte maturation inhibitor (Provot et al., 2006). Interestingly, previously reported *nkx3.2* mutant zebrafish display increased length of the proximal radials of the dorsal and anal fins (Waldmann et al., 2020), which is the opposite phenotypic outcome to what we saw in gdf5 mutant zebrafish fins.

The stimulation of Gdf5 in cultured human chondrocytes inhibits expression of the cartilage extracellular matrix degrading enzymes, including MMP13 (Enochson et al., 2014). In the pectoral fins of *gdf5* mutant zebrafish during the endoskeletal disc segmentation, when chondrocytes become mature and hypertrophic we didn’t observe chondrocytes in the most affected posterior radials, which could be the possible outcome if Gdf5 deficiency results in the increase of the MMP13 levels. However, in the median fins, we could not see any obvious differences in mature chondrocyte and cartilage extracellular matrix characteristics between wild-type and mutant fish. Furthermore, our analysis of adult *gdf5* mutant zebrafish revealed no difference in ossification process within pectoral and median fin skeletal elements when compared to wild-type fish. Further investigation is necessary in order to show the effect of *gdf5* inactivation on chondrocyte maturation, extracellular matrix organization, and mineralisation progress in zebrafish mutants and explain the mechanism for observed truncated skeletal elements.

Gdf5 has been implicated in the formation and positioning of joints in mouse and chicken autopods (Storm and Kingsley, 1996). It has been suggested to be involved in joint formation since it is expressed in joint forming regions in mouse autopods and Gdf5-deficient *brachypodism* mice display occasional joint defects (Storm and Kingsley, 1996). Several studies however argue against a direct role of *gdf5* in joint formation, but rather suggest the joint phenotype observed in *brachypodism* to be a secondary effect of the mutation (Francis-West et al., 1999b; Merino et al., 1999). This hypothesis is supported by studies showing a supply of exogenous Gdf5 is not sufficient to initiate ectopic joint formation but rather enhances cartilage growth differentiation (Merino et al., 1999; Storm and Kingsley, 1999).

We additionally analysed articulations in dorsal and anal fins between proximal and distal radials of adult gdf5 homozygote mutant and wild-type fish. *gdf5* is expressed in the distal part of the proximal radial and within the segmentation zone during distal radial development (Crotwell et al., 2001). Histological analysis revealed intact joints articulating proximal and distal radials, which indicates that Gdf5 loss does not affect correct segmentation. Interestingly, we could detect a thin layer of joint-specific cells showing similarities to articular cartilage in synovial joints. Furthermore, the cavity of those joints displayed the presence of a capsule, which could indicate a lubricin-filled cavity. To our knowledge there are currently no studies demonstrating the presence of lubricated synovial joints in median fins of zebrafish or other actinopterygian fishes. The presence of lubricin has been only demonstrated in jaw and pectoral fin joints of zebrafish pointing to their synovial joint-like properties (Askary et al., 2016). Whether and to what extent Gdf5 is involved in the formation of the primary jaw joint in zebrafish has not been investigated so far, however the expression of *gdf5* has been previously demonstrated in the jaw joint and the basihyal (Reed and Mortlock, 2010). Our analysis of zebrafish larvae at 9 dpf did not detect any skeletal phenotype in the primary jaw joint or in the craniofacial structures of *gdf5* mutants. We did however detect a small difference in size between the wild-type and homozygote mutant group at 9 dpf, possibly reflecting a slight delay in chondrogenesis of craniofacial structures, however this difference was not noticeable in juvenile or adult zebrafish (Supplemental Figure 1).

Interestingly, there is another member of GDF family colocalised in jaw joints of zebrafish, namely *gdf6a* (Reed and Mortlock, 2010). Inactivation of Gdf6 in mice leads to fusion of wrist and ankle joints and defects of middle ear joints articulating the middle ear ossicles (Settle et al., 2003). Since Meckel’s cartilage and the palatoquadrate in teleosts are homologous to the mammalian middle ear ossicles malleus and incus, *gdf6* knockout in zebrafish may lead to defects within first arch cartilage elements including the primary jaw joint. Mice deficient for both Gdf5 and Gdf6 display defects that do not occur in either single mutant comprising loss of bones and fused joints in limbs and the vertebrae (Settle et al., 2003). It is therefore interesting to speculate whether Gdf6 could also compensate for loss of *gdf5* during craniofacial development of zebrafish.

In summary our study for the first time demonstrates the importance of Gdf5 for the paired and median fin endoskeleton development in zebrafish and reveals that the severity of the effect increases from anterior to posterior side of the elements.

## 4 EXPERIMENTAL PROCEDURES

### 4.1 Ethical statement

All animal experimental procedures were approved by the local ethics committee for animal research in Uppsala, Sweden (permit number 5.8.18-18096/2019). All procedures for the experiments were performed in accordance with the animal welfare guidelines of the Swedish National Board for Laboratory Animals.

### 4.2 CRISPR/Cas9 target design

A sgRNA targeting the second exon of the zebrafish *gdf5* gene with no predicted off-target effects was designed using the online software CHOPCHOP (Labun et al., 2016), 5’ GAGAGTCCATGTGCTTCTTT 3’. The second base of the target was modified to “G” for T7 transcription without modifications. Preparation of the sgRNA was performed as previously described (Varshney et al., 2015), creating a fragment consisting of the T7 promotor, the targeted gene-specific sequence, and the guide core sequence. sgRNA synthesis was performed by in vitro transcription using the HiScribe T7 High Yield RNA Synthesis Kit (New England Biolabs, Ipswich, MA). Cas9 mRNA was prepared by in vitro transcription with the mMESSAGE mMACHINE T3 Transcription Kit (Life Technologies, Carlsbad, CA) using 500 ng of linearised plasmid that was retrieved from 5 μg of p-T3TS-nCas9n plasmid (plasmid #46757; Addgene, Cambridge, MA) digested with XbaI (New England Biolabs, Ipswich, MA). The products were purified, and their integrity was assessed using a denaturation gel.

### 4.3 Generation of zebrafish mutant line

Fertilised zebrafish (*Danio rerio*) eggs were obtained by natural spawning of wild-type AB line. Embryos were injected at the one-cell stage with 150 pg of Cas9 mRNA and 50 pg of each sgRNA in RNase-free water as previously described (Varshney et al., 2015), and maintained at 28.5°C in E3 medium (Westerfield, 2000). The efficiency of the target was estimated by the CRISPR-Somatic Tissue Activity Test (STAT) methodology in eight embryos at two days post-injection, as previously described (Carrington et al., 2015). The injected founder zebrafish (F0) were raised and incrossed. For genotyping the F1 zebrafish, DNA was extracted from a 1-3 mm amputation of the adult zebrafish caudal fin by lysing the tissue in 30 μl of 50 mM NaOH for 20 min at 95°C, adding 60 μl of 0.1 mM Tris and diluting the obtained material (1:10). For the initial genotyping step, FLA analysis was used. 2 μl of DNA (50-200 ng) was added to Platinum Taq DNA Polymerase. The PCR mix was incubated at 94°C for 12 min followed by 35 cycles of 94°C for 30 sec, 57°C for 30 sec, 72°C for 30 sec, and final extension at 72°C for 10 min. Size determination was carried out on a 3130XL ABI Genetic Analyzer (Applied Biosystems, Waltham, MA) and the data were analysed using the Peak Scanner Software (Thermo Fisher Scientific, Waltham, MA). For the fish that screened positive for the variant, the FLA results were confirmed by Sanger sequencing (forward primer with m13 tag: 5’-TGTAAAACGACGGCCAGTACGTCTTCAACATCAGCTCA-3’; Reverse primer with PIGtail: 5’-GTGTCTTCCAGATGTCAAACACCTCCC-3’). One strain with an allele containing a frameshift deletion resulting in a premature stop codon (*gdf5*^uu3703^) was selected for further experiments. The identified F1 founders were crossed with wild-type zebrafish (AB strain), and their adult offspring (F2) were genotyped. Heterozygous F2 fish of mutant line *gdf5*^+/uu3703^ were incrossed and the offspring were screened for homozygous mutants.

### 4.4 Skeletal staining

Zebrafish wild-type and mutant fish were euthanised, fixed in 4% PFA and transferred to 50% ethanol. For OPT analysis, 9 dpf larvae were genotyped prior to double cartilage and bone staining by clipping a small piece of the tail and performing genotyping by FLA as described above. Staining of cartilage and bone was done based on the previously published protocol by Walker and Kimmel (2007). For double staining of cartilage and bone, specimens were immersed in double staining solution of 99% alcian blue solution (0.02% Alcian Blue 8 GX, 50mM MgCL2, 70% ethanol) and 1% alizarin red solution (0.5% Alizarin Red S). After staining overnight, specimens were washed twice with 50% ethanol and then immersed in water for 2 hours before being bleached in a solution of 1.5% H202 and 1% KOH until pigmentation was removed. Specimens 30 dpf and older were then immersed in trypsin solution (1% trypsin, 35% sodium tetraborate) for 30 minutes followed by incubation in a solution of 10% glycerol and 0.5% KOH for 1 hour. All specimens were imaged with a Leica M205 FCA microscope in a solution of 50% glycerol and 0.25% KOH, followed by storage in 50% glycerol and 0.1% KOH.

### 4.5 Optical Projection Tomography

A custom-built Optical Projection Tomography (OPT) system was used for imaging of the zebrafish embryos fixed at 9 dpf and stained cartilage and bone (Sharpe et al., 2002; Zhang et al., 2020). The OPT system, reconstruction algorithms, and alignment workflow were based on the previously described method (Allalou et al., 2017). All embryos were kept in 99% glycerol before they were loaded into the system for imaging. The rotational images were acquired using a 3x telecentric objective with a pixel resolution of 1.15 μm/pixel. The tomographic 3D reconstruction was done using a filtered back projection (FBP) algorithm in MATLAB (Release R2015b; MathWorks, Natick, MA) together with the ASTRA Toolbox (Palenstijn et al., 2013). For the data alignment, the registration toolbox elastix (Klein et al., 2010; Shamonin et al., 2014) was used. To reduce the computational time all 3D volumes in the registration were down-sampled to half the resolution.

The registration workflow was similar to the methods described by where the wild-type fish were initially aligned and used to create an average reference fish using an Iterative Shape Averaging (ISA) algorithm(Rohlfing et al., 2001). All wild-type (*n*=18) and *gdf5*^uu3703/uu3703^ (*n*=20) zebrafish were then aligned to the reference. No voxel-wise method was used after alignment to detect voxels that are significantly different between the groups due to big size variation between individuals.

### 4.6 Histological analysis

Zebrafish at 60 dpf were fixed in 4% PFA at 4°C overnight. Decalcification was performed in 20% EDTA solution for 7 days Following decalcification, the samples were dehydrated through a series of ethanol washes in 2x 70% ethanol for 45 minutes each, 2x 90% ethanol for 45 minutes each, 95% ethanol for 1 hour and 2x 99.5% ethanol for 45 minutes each. Ethanol was replaced by xylene and samples were incubated for 10 minutes followed by a 5 minute incubation in fresh xylene. The xylene was replaced by melted Paraffin and samples were incubated at 65°C overnight. The following day, samples were incubated in freshly melted Paraffin at 65°C prior to embedding. Embedded fish were sectioned at a thickness of 6 μm. For Hematoxylin and Eosin (HE) staining, 6 μm paraffin sections were deparaffinised in xylene (2x 10 min) and re-hydrated through a series of ethanol washes (2x 99.5%, 2x 95%, 90%, and 70% for 10 minutes each) to distilled water. Sections were subsequently stained in Hematoxylin for 7 minutes followed by a 15 minute wash under running water. Sections were dipped 2x in 70% ethanol follow by 90 second staining in Eosin. After the staining, sections were dipped in a series of ethanol (2x 95% ethanol, 1x 99.5% ethanol, then 5 minutes in 99.5% ethanol) followed by 2 dips in xylene prior to mounting with Cytoseal. All slides were imaged with 40x objective on Hamamatsu NanoZoomer S60 Digital Slide Scanner.

### 4.7 Micro-computed tomography and segmentation

Three wild-type and five *gdf5*^uu3703/uu3703^ zebrafish were fixed at 90 dpf and analysed with micro-Computed Tomography (μCT, SkyScan 1172, Bruker microCT, Belgium) at a voltage of 60 kV, a current of 167 μA, and an isotropic voxel size of 2.44 μm. The specimens were placed in 2 mL Eppendorf tubes filled with 1% agarose. Cross-sections were reconstructed using software package NRecon (NRecon 1.6.10, Bruker microCT, Belgium). When working with the whole-body scan data, image stacks were downsampled 2x to a 4.88 μm voxel size in Fiji (Schindelin et al., 2012). BMP image stacks obtained by μCT were imported into, segmented, and imaged using VGStudio MAX version 3.3.2 (Volume Graphics, Germany).

## ACKNOWLEDGEMENTS

TH was supported by Vetenskapsrådet (Starting grant 621-2012-4673). The development of the OPT system was funded by a development project at SciLifeLab Uppsala (2017) and a Technology Development grant at SciLifeLab (2018) both awarded to AA. VG Studio MAX licence and some lab expenses were covered by a Wallenberg Scholarship awarded to Prof. Per E. Ahlberg, who also offered thoughtful comments on the manuscript. We thank the Genome Engineering Zebrafish facility in SciLifeLab Uppsala for generating CRISPR/Cas9 mutant, and Prof. Åsa Mackenzie for access to the Hamamatsu NanoZoomer S60 Digital Slide Scanner.

## DECLARATION OF INTEREST

The authors declare no conflicts of interest.

## AUTHOR CONTRIBUTIONS

TH, LW, and JL designed this project; all authors performed experimental work and analysed the data; TH, LW, and JL wrote the paper with contributions from other authors.

## SUPPLEMENTARY MATERIAL

**SUPPLEMENTAL FIGURE 1.**
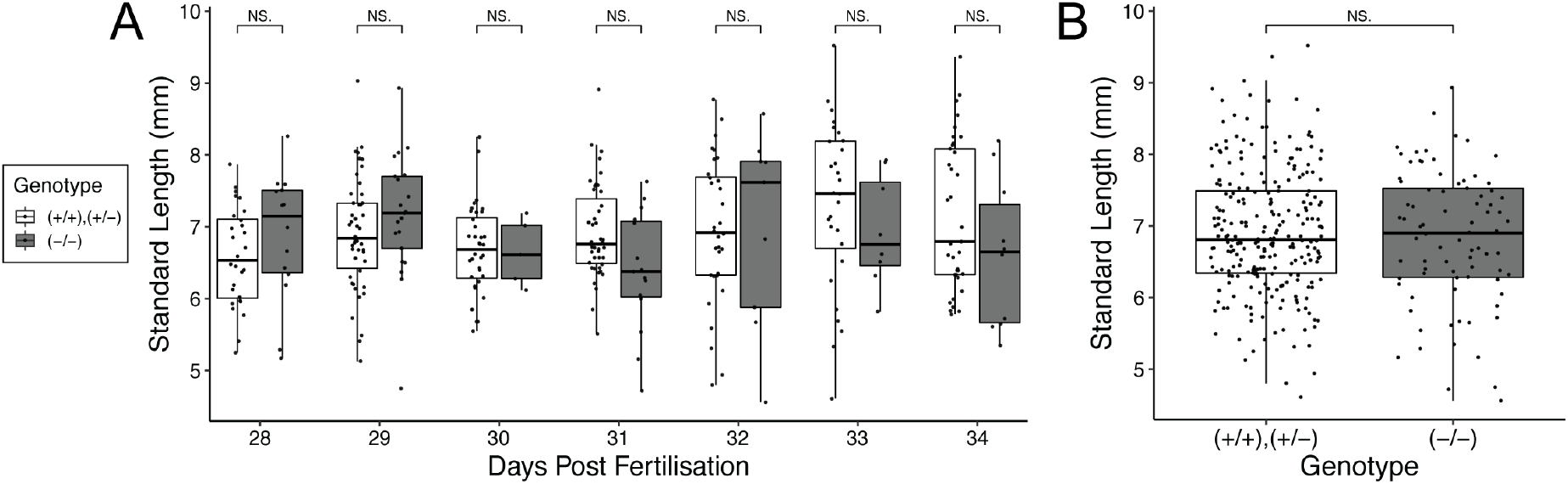
No significant standard length differences between juvenile wild-type and *gdf5^uu3703/uu3703^* zebrafish. **(A)** Standard lengths of sampled zebrafish between 28-34 dpf, broken down by age. **(B)** Pooled comparison of the standard lengths of all juvenile zebrafish in **(A)**. N=332. Genotype was inferred from the presence or absence of a skeletal phenotype. All wild-type to gdf5 mutant comparisons found no significant difference (P>0.050, Wilcoxon test), with the exception of 31dpf which was significant in isolation (P=0.024), but not after P-value adjustment for false discovery rate (Benjamini and Hochberg, 1995).

**SUPPLEMENTAL VIDEO 1 | Optical projection tomography of wild-type and *gdf5*^uu3703/uu3703^ zebrafish at 9dpf. (A, B)** Examples of raw rotational imaging of single 9 dpf zebrafish heads captured during OPT. **(C, D)** Rotating summary projection of 3D visualization of skeletal structures produced by processing the rotational imaging of single fish. **(E, F)** as **(C, D)** but averaging the results from all fish. **(G, H)** As **(E, F)** but maximum projection view.

## Notes

### Competing Interest Statement

The authors have declared no competing interest.

